# Wavefront estimation through structured detection in laser scanning microscopy

**DOI:** 10.1101/2024.12.31.630574

**Authors:** Francesco Fersini, Alessandro Zunino, Pietro Morerio, Francesca Baldini, Martin J. Booth, Alessio Del Bue, Giuseppe Vicidomini

**Author notes:** OncoBreast, Inserm - Centre de recherches en cancérologie de Toulouse, Toulouse, France (Francesco Fersini).

## Abstract

Laser scanning microscopy (LSM) is the base of numerous advanced imaging techniques, including confocal laser scanning microscopy (CLSM), a widely used tool in life sciences research. However, its effective resolution is often compromised by optical aberrations, a common challenge in all optical systems. While adaptive optics (AO) can correct these aberrations, current methods face significant limitations: Aberration estimation, which is central to any AO approach, typically requires specialized hardware or prolonged sample exposure, rendering these methods sample-invasive, and less user-friendly. In this study, we introduce a simple and efficient AO approach for CLSM systems equipped with a detector array – the same of super-resolved image-scanning microscopy – and an AO element for beam shaping. We demonstrate that imaging datasets acquired with a detector array inherently encode aberration information. Leveraging this property, we developed a custom convolutional neural network capable of decoding aberrations, up to the 11^*th*^ Zernike coefficient, directly from a single acquisition. This method enables a new generation of AO implementations for LSM, offering an accessible solution that minimizes sample stress while achieving high-resolution, and aberration-free imaging.

## Introduction

Confocal laser-scanning microscopy (CLSM) is a pivotal imaging tool in life science research (1–4). While its beam-scanning architecture results in lower imaging temporal resolution (frames *per* second) compared to its wide-field counterparts, this design also introduces significant advantages: By employing a physical pinhole to eliminate out-of-focus fluorescence signals, CLSM provides exceptional optical sectioning (5), making it ideal for three-dimensional imaging of thick samples. Additionally, CLSM can be combined with spectroscopy (hyperspectral imaging) to analyze the fluorescence emission spectrum and/or fluorescence lifetime of the sample (6). This integration allows researchers to correlate the sample’s structural features with its functional properties and phenotyping (7).

However, like all optical microscopy techniques, CLSM is susceptible to system- or specimen-induced aberrations, which can distort the optical wavefronts of both the illumination and detection light, significantly degrading image quality. Adaptive optics (AO) systems address this issue by employing reconfigurable elements, such as deformable mirrors (DMs), spatial light modulators (SLMs), or adaptive lenses, to dynamically adjust the optical wavefronts and effectively compensate for aberrations (8–16). A key element for the success of an adaptive optics approach is its ability to detect optical aberrations using simple and non-invasive – both for the optical architecture and the sample – methods. There are two primary classes of aberration detection methods: One involves direct measurement of aberrations using wavefront sensors, while the other infers aberration directly from the images, hence the name “sensorless”. In wavefront sensor methods, phase aberrations are measured directly using a Shack-Hartmann sensor or an interferometer, offering speed in the estimation of the aberration but suffering from complexity in optical design and non-common path errors (17– 19). Conversely, sensorless methods utilize simpler optical designs but depend on iterative estimation of aberrations by evaluating specific image quality metrics, such as total signal intensity or spatial frequency-based sharpness, in response to phase modulations of the optical wavefront (20–23). This leads to time-consuming processes with repeated sample exposures, potentially causing photo-damage or motion-related errors.

In recent years, sensorless AO methods driven by machine learning (ML) have emerged as efficient alternatives to traditional techniques. These approaches significantly reduce the number of sample exposures (i.e., phase modulations) required for aberration correction, offering gentler solutions for specimens while maintaining robustness. Convolutional neural networks (CNNs), in particular, are powerful tools for automatically extracting relevant features directly from raw data without requiring manual intervention. Additionally, CNNs are capable of handling large and complex datasets and can be trained on simulated data, overcoming the challenge of limited experimental data availability.

Early implementations of these methods relied on access to the system’s point spread function (PSF), either directly or through imaging point-like structures, such as fluorescent beads (24–27). However, this dependence on PSFs limited their applicability to a broader range of imaging scenarios. More recently, advances in machine learning have enhanced the efficiency and adaptability of neural network-based AO methods by removing the need for point-like source imaging. For example, transfer learning frameworks and novel neural network architectures that integrate physical principles of microscope image formation directly into their models have shown promising results (28, 29). Despite these improvements, some degree of phase modulation is still required, and there remains significant potential to further enhance efficiency by providing more informative datasets for neural network training.

Here, we demonstrate how sensorless AO for CLSM can improve the versatility of aberration estimation with a sample-friendly approach by leveraging an emerging class of fast detector arrays. Unlike conventional single-element detectors, such as photomultiplier tubes, these array detectors capture spatial information that is otherwise lost, facilitating the access to the optical aberrations distorting the images. This unique information can be harnessed by a NN-based sensorless approach, requiring only a single imaging experiment to detect most aberrations. Notably, these arrays detector are the same ones emerging as the gold standard for image-scanning microscopy (ISM), which is gradually replacing CLSM. ISM enables effective super-resolved imaging while retaining all the functionalities of traditional confocal microscopy. In this context, we demonstrate that our approach introduces a novel CLSM implementation that synergistically and straightforwardly combines the advantages of AO and ISM. We named this approach image scanning microscopy with adaptive optics (ISMAO).

Image scanning microscopy (30–33) encompasses a class of techniques which achieves effective super-resolution by leveraging the theoretical ability of confocal microscopy to double the resolution of conventional microscopy. In confocal microscopy, reducing the size of the pinhole decreases the point spread function (PSF), potentially improving resolution. However, this approach also significantly reduces the signal from the focal point, deteriorating the image signal-to-noise ratio (SNR) and negating the resolution gain. In laser-scanning architectures, such as the CLSM, ISM overcomes this limitation by substituting the pinhole and single-element detector with a fast, pixelated detector. Examples of such detectors include the AiryScan (34) and asynchronous read-out single-photon avalanche diode (SPAD) arrays (35, 36). These detectors capture microimages for each focal region probed by the laser during raster scanning, preserving spatial information that would otherwise be discarded. The captured microimages are processed to produce a final image with effectively doubled spatial resolution and improved signal-to-noise ratio (SNR). Specifically, by treating each element of the pixellated detector as a small pinhole, the microimages recorded during the scanning are transformed into a series of CLSM images (referred to as scanned images), but overall the detector array collects all fluorescence photons. By means of a simple reconstruction algorithm named pixel-reassignment (37–39) the scanned image are fused in a final high-resolution and high-SNR image. Essentially, the scanned images are spatially registered, by means of a phase-correlation approach, before integration. More recently, novel ISM reconstruction algorithms – in the context of pixel assignment (40) or multi-image deconvolution (41)– have been introduced, enabling resolution enhancement without compromising the optical sectioning capability of CLSM. In this work, we demonstrate that the ISM dataset inherently contains information not only to enhance resolution but also to extract the optical wavefront of the illumination and detection beam. The array detector functions similarly to a scanning Shack-Hartmann sensor, enabling the decoding of the aberrations without the need for iterative approaches or dedicated sensors. In other words, the same detector used for imaging also performs wavefront sensing. To this end, we developed a pipeline to pre-process the data in order to create a specimen-free dataset for a custom CNN. This latter decodes the phase information from the data, returning the coefficient of the Zernike polynomials required to describe the wave-front error that corrupts the images. We trained the CNN on synthetic datasets generated through a comprehensive image formation simulation framework, parametrizing a variety of imaging conditions. We improve this approach by fine-tuning the network with just a limited amount of experimental data, thanks to the transfer learning approach which enhances the network’s performance in practical applications. Finally, we demonstrated the capability of ISMAO to correct aberrations in a custom-built ISM system featuring a SPAD array detector and a DM integrated into both the excitation and detection paths.Our results represent a step forward toward a deeper comprehension of the physics of structured detection and its potentialities for applications, such as imaging in thick specimens.

## Results

### Principle of ISMAO: aberrations encoding

In this section, we demonstrate that ISM imaging datasets inherently encode information about the optical aberrations present in the system, which distort the wavefronts of both the illumination and detection light paths.

In an ISM setup (Fig.1a, Suppl. Fig. S1), a focused laser beam excites the sample point-by-point at the scan coordinates x_s_ = (*x*_s_, *y*_s_). Then, the microscope collects and descans the fluorescence light, imaging the probed region onto the sensitive area of the detector array and generating a microimage with detector coordinates x_d_ = (*x*_d_, *y*_d_). Each pixel of the detector array acts as a small pinhole, located in a unique position of the detector plane. With a full scan, the signal collected by each pixel arranged along the scan co-ordinates forms a confocal-like image of the same specimen observed from a slightly different point-of-view (42). Thus, the ISM dataset *i*(*x*_*s* |_ *x*_*d*_) is four dimensional and can be described either as a large collection of microimages (Fig. 1b) or a as a small collection of scanned images (Fig. 1c). The latter perspective allows for the following description of the ISM image formation:

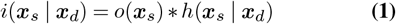

where *o*(*x*_*s*_) is the object function – namely the distribution of fluorophores in the specimen – and *h*(*x*_*s*|_ *x*_*d*_) is the point spread function (PSF) of the imaging system – given by the product of *h*_exc_(*x*_*s*_) with *h*_det_(*x*_*s*|_ *x*_*d*_), the excitation and detection PSFs. The first is the focused excitation beam which is always aligned to the optical axis, while the second is the emission PSF shifted to the position *x*_*d*_ of the detector element that generated the image. In an ideal scenario, both excitation and detection PSFs are diffraction-limited. Thus, their product is narrower than the diffraction limit, enabling super-resolution. However, in a realistic scenario, the optical performances might be compromised by wavefront distortions, which degrade the imaging quality and limit the effective image resolution and signal-to-noise ratio (SNR). The specimen itself is the most common source of optical aberrations. As light travels through the layers of a thick sample (Fig. 1d), it accumulates wavefront distortions, causing both the excitation and fluorescence light to be affected by aberrations. The wavefront error at the objective pupil plane is directly reflected in the shape of the PSF, which is uniquely distorted by various optical aberrations. Consequently, conventional wavefront sensing methods —-such as through-focus bead fitting – require access to the system PSF, typically obtained by imaging point-like structures (43). In contrast, we demonstrate that wavefront information is inherently encoded in the raw ISM dataset, enabled by the unique array detection method employed in ISM. This innovation eliminates the need for imaging point-like structures for wave-front estimation. Indeed, all scanned images have the same source and represent the same specimen. Therefore, any difference between these images has to be ascribed to different PSFs. Since the structure of the PSFs uniquely encodes aberrations, we can leverage the diversity of the images to recover the wavefront without the need for a dedicated sensor other than the detector array itself. Nonetheless, we need a strategy to decode the wavefront information independently from the observed specimen. To this end, we need to measure a quantity that depends uniquely on the PSFs of the system. A natural choice would be the fingerprint *f* (*x*_*d*_), which is given by the sum of all the micro-images collected during the scan (Fig. 2a). Assuming that the signal is largely in focus, the fingerprint is independent of the structure of the specimen and depends uniquely on the cross-correlation between *h*_*exc*_ and *h*_*det*_. In other words, it represents the photon detection probability on the detector plane.

**Fig. 1.**
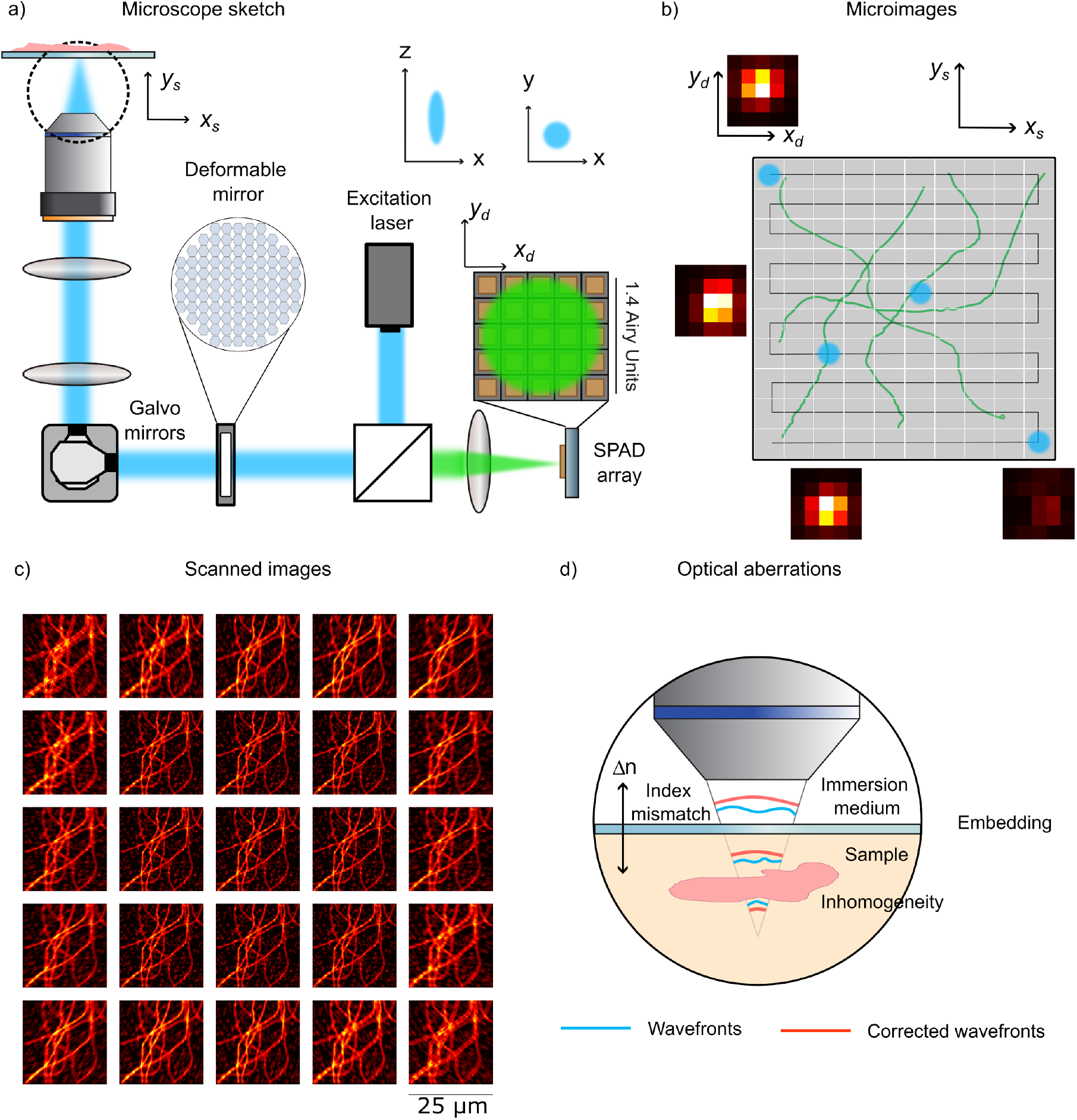
a) Sketch of the ISMAO setup. The system is equipped with a detector array in the detection arm and an Alpao deformable mirror with 97 actuators placed in a conjugate plane with the objective lens. The multidimensional output of ISMAO is composed of b) microimages - generated during the raster scanning of the sample – where for each scanned point, a 5 × 5 intensity image is obtained and c) the 25 confocal-like scanned images collected by each element of the detector array. d) An example of wavefront deformation caused by a thick sample with a refractive index mismatch, which generates an ill-defined diffraction-limited excitation spot.

**Fig. 2.**
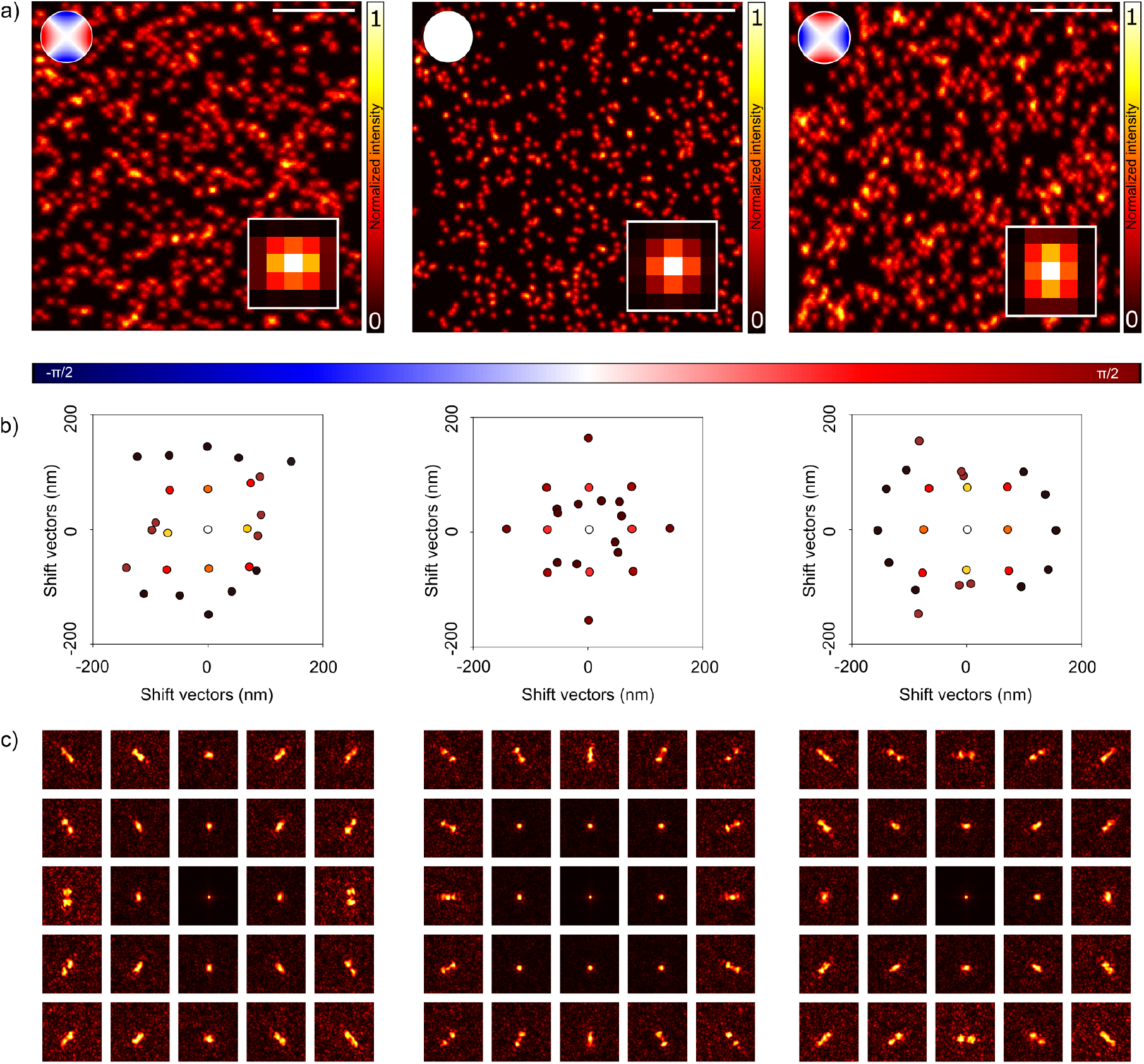
This figure demonstrates how an array detector inherently encodes optical aberration information. a) The sum of scanned images is displayed, with phase modulation introduced by a deformable mirror. The phase mask used in each simulation is indicated in the top-left corner. Aberrations in the PSF alter the intensity distribution across the active area of the sensor, modifying the fingerprint shown in the bottom-right corner of each simulation. Scale bar 10 *µ*m. b) Shift vectors provide an alternative method for estimating optical aberrations. These vectors are calculated from the intensity values corresponding to the maximum correlation between the central image and others. While less comprehensive than fingerprints, they offer a direct measure of aberrations. Each sensor element is assigned a shift vector, represented using the same color coding as the fingerprint. c) Correlograms encode optical aberration information directly and are independent of the sample structure. Unlike scanned images, correlograms preserve the physical and optical properties of the PSF, making them a more robust and generalizable approach for aberration analysis compared to shift vectors.

Through simulations, we illustrate how aberrations alter the intensity distribution of the fingerprint, becoming no longer sharply centered in the detector array but spread across the entire detector plane (Fig. 2a, and Suppl. Figs. S3, S4, S5, S6). Therefore, aberrations directly affect the structure of the fingerprint distribution, enabling almost direct access to the PSF of the microscope of the system without the need for point-like sources – acting as guide-stars. Nonetheless, to maintain a temporal resolution compatible to fast scanning the ISM detector array posses only a few sensitive elements (twenty-five for the SPAD array detector used in this work), which result in an inadequate sampling to robustly retrieve a complex wavefront. A second approach for quantifying the aberrations lies in the measurement of the shift vectors, namely the translations required to maximize the similarities between the raw images of the ISM dataset to the one generated by the central element of the detector array. Indeed, the scanned images are approximately identical, but shifted and rescaled, as long the magnification of the system is sufficiently high (39). A well-established algorithm, known as adaptive pixel reassignment (APR), exploits this concept to shift back and sum the raw images, reconstructing a single super-resolution image from the raw data (44, 45). Given the linearity of fluorescence image formation, a shift in the images uniquely relates to a shift in the PSFs. Therefore, if one or multiple aberrations distort the shape of the PSFs, shift vectors change accordingly, encoding the wavefront information in their pattern (Fig. 2b). However, shift-vectors are well-defined only for single-peaked PSFs, a condition fulfilled only without strong aberrations and using large magnification values. Given the limitations of the previous two approaches, we developed a more comprehensive strategy which inherently contains all the information provided by the two approaches. In detail, we leverage the phase cross correlation between the scanned images to calculate the correlograms, which depend uniquely on the PSFs of the microscope (Fig. 2c). The cross-correlation between an image of the ISM dataset and the reference image at the center of the array detector (*x*_*d*_ = 0) is the following:

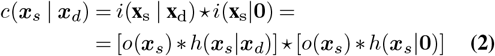

where * and * are the convolution and the cross-correlation operator, respectively. Then, we apply the Fourier transformation with respect to the scan coordinates *x*_*s*_. Using the convolution theorem, we obtain

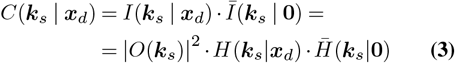

where the overline represents the complex conjugate operation, *k*_*s*_ is the scan spatial frequency, and the uppercase letters indicate the Fourier transform of the quantities previously indicated with the lowercase letter. To discard the object contribution, we need to normalize the correlation in Fourier space.

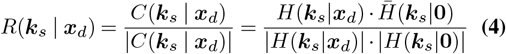

Finally, we obtain the correlograms by inverting the Fourier transform and we normalize with the fingerprint

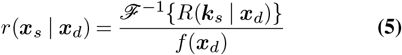

The result depends uniquely on the PSFs of the microscope regardless of the structure of the specimen. Notably, the coordinates of the maximum of the correlegrams are, by definition, the shift vectors. Therefore, we designed a quantity that encodes the highest amount of information about the wave-front without the bias of a specific sample.

### Principle of ISMAO: aberrations decoding

In this section, we demonstrate how the optical aberrations encoded within the ISM dataset can be effectively decoded and used to control an AO element for precise aberration correction. Two approaches can be envisioned for this task. The first consists of inverting the correlogram formation model by solving a minimization problem of the error between the model and the data. This approach leverages an explicit physical model but requires a relatively slow and computationally intensive minimization procedure. The second approach uses a CNN to retrieve the coefficient of the Zernike polynomials from the data. This method does not require a model, which is learned by the network itself through the training procedure. We performed this latter using physically accurate numerical simulations of correlograms. Furthermore, the DM in our custom setup also enables a straightforward acquisition of experimental data which can be used to refine further the network. Given the ease of generation of data for the training and the speed of the CNN predictions, we opted for the CNN approach. Indeed, the speed of a trained neural network better fits the needs of real-time adaptive optics systems, where aberrations must be estimated rapidly to drive the corrective optical elements.

We implemented two closely related CNNs that differ mainly in their input datasets. The first CNN (CNNx1) receives as input the normalized correlograms obtained from a single focal plane ISM acquisition. In contrast, the second CNN (CNNx3) processes normalized correlograms derived from a three-planes ISM acquisition. To acquire more information, we introduce a defocus as large as half the depth-of-field, DoF = (2λ*n*)*/*(NA^2^). For CNNx3, two additional focal planes, positioned at DoF*/*2 and +DoF*/*2 relative to the original plane, are included as inputs, where DoF represents the depth of field of the optical system. This approach utilizes multi-plane data to improve aberration decoding, particularly for aberrations that disrupt axial symmetry, such as astigmatism or spherical aberrations.

To generate the training set, we used the open-source BrightEyes-ISM Python package (46).To generate ISM images under varying optical aberrations, we parameterize the wavefront distortions at the objective pupil plane – both for the excitation and emission PSFs – using a Zernike polynomial basis:

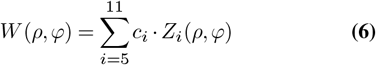

Here, *ρ* is the normalized radius of the pupil, *φ* is the azimuthal angle of the pupil, *c*_*i*_ is an amplitude expressed in radians, and *Z*_*i*_ is the Zernike polynomial *i* (using the Noll index notation). Note that we neglect the first four orders of aberrations (piston, tip, tilt, and defocus) because they describe a simple misalignment that we are not interested in correcting through adaptive optics. Furthermore, we interrupt the series expansion at the 11^th^ polynomial to take into account only the primary aberrations. Neglecting higher-order aberrations balances computational efficiency while still addressing the most common aberrations encountered in microscopy. Once the ISM dataset is generated, we calculate the normalized correlograms to feed the CNN (Eq. 5). To account for experimental uncertainties during CNN training, we introduced several variations. The number of simultaneously occurring aberrations was randomly chosen, with their amplitudes sampled from a distribution defined as the sum of two Gaussian, each with *µ* = ±0.75, rad and *σ* = 0.3, and restricted to an RMSE range of ±*π/*2, rad (Fig. 3a). Additionally, the emission photon flux and pixel dwell time were adjusted to reflect diverse signal-to-noise imaging conditions (Fig. 3b-c). Variations in the numerical aperture (NA) of the objective were also introduced to simulate additional experimental conditions.

**Fig. 3.**
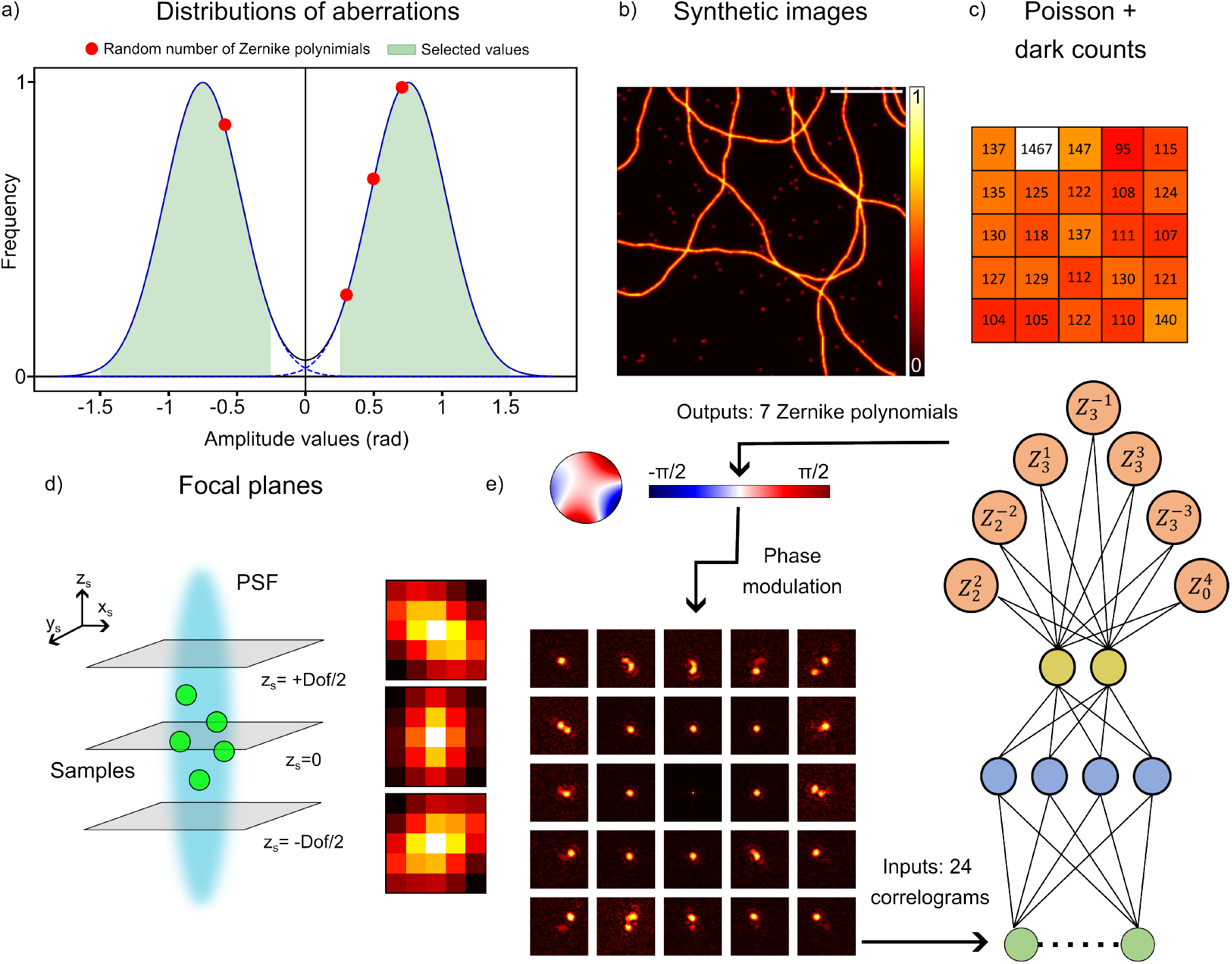
a) The first panel illustrates the density distribution of optical aberrations randomly selected to generate the simulated phase mask of the PSF. The green area under the curves represents the density distribution range considered for the simulations. The number of simultaneous aberrations is defined randomly; an example in the graph shows four selected aberrations. b-c) Show an example of a simulated phantom, including 3D tubulin filaments and beads, which are used for testing and the photon dark count collected during a 1-second acquisition with the array detector, replicating realistic detector noise during pixel dwell time acquisition. d) The fourth panel represents the different focal planes simulated. The axial asymmetry of certain aberrations highlights the advantage of collecting data from multiple focal planes to improve aberration identification. e) ISMAO approach. The correlograms (normalized with the fingerprint) are used as input data for the CNN, which is based on the AlexNet structure. It has seven outputs corresponding to seven Zernike polynomial coefficients from 5^*th*^ to 11^*th*^. Alternatively, the CNN can process 3 × 24 correlograms if data is collected from three focal planes. The outputs are then used to drive an adaptive element in the microscope to apply phase modulation to the experimental PSF and compensate for the predicted aberrations.

### ISMAO validation on synthetic images

To evaluate the performance of the ISMAO approach, we implemented and tested two CNNs models. Both models were trained on the same synthetic dataset to ensure consistency and reproducibility. The validation protocol was conducted in two phases.

In the first phase, the CNNs were employed to identify individual aberrations, with only one aberration introduced at a time. This phase aimed to assess the models baseline capability to predict aberrations individually. For this test, the validation dataset consisted of 490 simulations, corresponding to 7 aberration types, each evaluated at 7 different amplitude points, with 10 repetitions for each aberration amplitude. Specifically, the amplitude of the test dataset ranged from -1.5 to 1.5 rad, with a step size of 0.5 rad. This experiment demonstrated that CNNx3 more accurately tracked the trends of imposed aberrations compared to CNNx1 (Fig. 4a, and Suppl. Fig. S7). While CNNx1 provided predictions comparable to CNNx3 for smaller aberrations, however, its performance suffered for larger aberrations. Additionally, CNNx3 exhibited enhanced robustness in handling aberrations that disrupted axial symmetry, such as astigmatism and spherical aberration. This trend was further confirmed by the correlation matrix (Fig. 4b), which evaluates the models ability to accurately identify specific aberrations. The correlation matrix reflects the strength of the relationship between real and predicted data across the set of aberration variables.

**Fig. 4.**
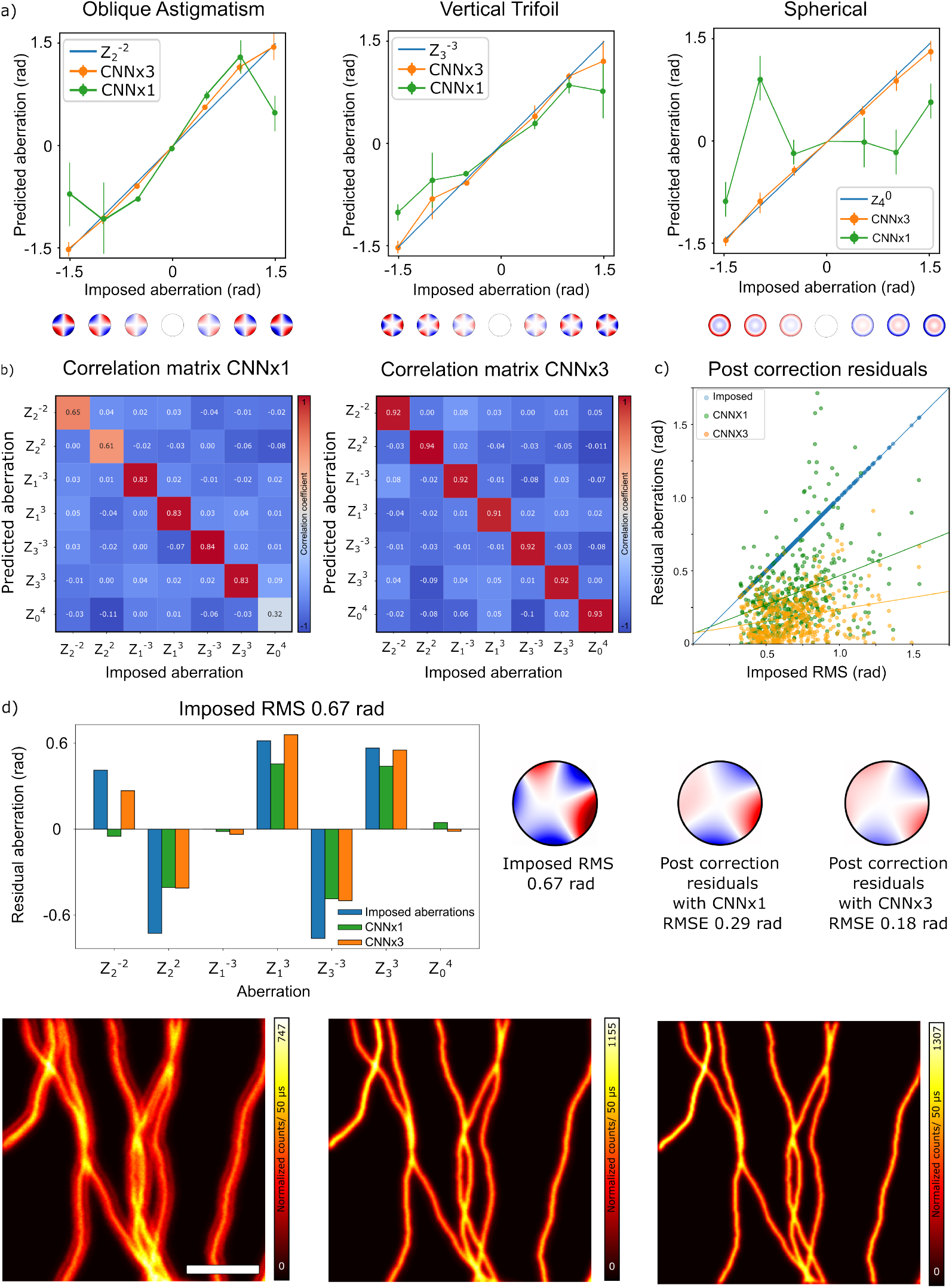
a) Comparative analysis of phase retrieval using two CNNs. In this analysis, a single aberration is considered at a time, focusing on oblique astigmatism 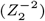 and vertical trefoil 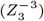 and spherical 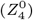 aberration, within the range of ([−1.5; 1.5]) rad, with an interval of 0.5 rad. The plot shows that CNNx3 (orange) better approximates the imposed aberration (blue) compared to CNNx1 (green). b) Correlation maps for both CNNs. CNNx3 achieves a correlation score of 92%, outperforming CNNx1, particularly in estimating aberrations with axial asymmetry. c) shows residual errors from 700 simulations after phase retrieval with CNNx1 and CNNx3. In order to visualize the error of the estimated aberration in a more complex scenario (where multiple and random number of aberration occur simultaneusly) d) provides a histogram of multiple aberrations with an RMSE value of 0.67 rad. The CNNx3 outperforms CNNx1, improving image quality and resolution after correction. Scale bar: 5 *µ*m.

In the second phase, a more complex scenario was introduced, wherein multiple aberrations were simultaneously imposed. This phase allowed for a comprehensive evaluation of each CNN ability to manage the compounded effects of overlapping aberrations, providing an assessment of their robustness and generalizability under more realistic conditions.

For this phase, we employed simulated datasets designed to replicate realistic experimental scenarios, where multiple aberrations commonly occur due to factors such as sample thickness, refractive index mismatches, and system-induced aberrations. To simulate this complexity, we generated a dataset of 700 simulations for each focal plane, with the number and amplitude of simultaneously occurring aberrations chosen randomly based on a double Gaussian distribution, as during CNN training.

We summarize the results for both CNN models using a scatter plot of the imposed RMS aberration versus the residual RMS error (Fig. 4c). The RMSE is computed as the difference between the imposed aberration and the phase retrieved by CNNx1 and CNNx3. The imposed RMS values ranged from 0.25 to 1.5 rad (values below 0.25 rad were discarded due to their negligible effect on image quality). The scatter plot clearly illustrates that CNNx3 consistently yielded lower residual errors compared to CNNx1.

To further assess model performance under challenging conditions, we conducted a synthetic experiment (Fig. 4d). In this scenario, a total RMS of 0.67 rad was imposed across multiple aberrations. Both CNN models successfully retrieved the imposed aberrations and applied corrections to the same images. Despite the complexity of this scenario, in which five aberrations with varying amplitudes were simultaneously present, both models were able to estimate the imposed aberrations. Notably, predictions from CNNx3 closely aligned with the ground truth, as shown in the histogram of imposed aberration values. Although CNNx1 showed good overall predictive accuracy, it exhibited limitations in specific cases, such as astigmatism, where its predictions were less precise compared to CNNx3, as expected.

Following the application of corrections derived from the predictions, CNNx1 reduced the RMSE to 0.29 rad, while CNNx3 achieved a more precise correction, with an RMSE of 0.18 rad. The corrected images highlight the robustness of the approach, demonstrating its ability to predict and compensate for aberrations in a single acquisition. Additionally, we observed that CNNx1 performed well in less complex scenarios but faced challenges when dealing with aberrations exhibiting significant axial asymmetry, such as the simultaneous presence of astigmatism and spherical aberration (Supp. Fig. S7). These results underscore the superior accuracy of CNNx3 in managing complex aberration profiles, while also demonstrating the efficiency and reliability of CNNx1 in less demanding scenarios, where a single acquisition can provide sufficient precision for aberration estimation.

### ISMAO validation on experimental images

To validate the ISMAO method on real datasets, we implemented a custom ISM system incorporating a 5 × 5 single-photon avalanche diode (SPAD) array detector and a deformable mirror (DM) positioned within the shared path of the excitation and emission beams. In this work, the DM serves a dual role: it can correct specimen- and system-induced aberrations identified through an AO method, such as our ISMAO approach, but it can also be used to introduce well-defined, controlled optical aberrations into the system. This latter capability enabled the implementation of a transfer learning approach, enriching the CNN training process with conditions not fully captured in purely synthetic datasets. Specifically, we collected a series of ISM datasets by applying specific single-coefficient aberrations. To expand the dataset size, we employed data augmentation strategies and integrated this relatively small real dataset (5–10% of the total training dataset) with the synthetic dataset. The combined synthetic and real dataset was then used to train the CNNs effectively. For transfer learning, we used images of lipid droplets (47, 48) (Fig. 5a) and fluorescent beads (Supp. Fig. S8). Specifically, we captured a series of ISM datasets by introducing single aberrations at varying strengths using the deformable mirror. This experimental data was incorporated into the dataset for fine-tuning the model, improving its ability to detect and correct optical aberrations in the imaging scenarios of our custom microscope.

**Fig. 5.**
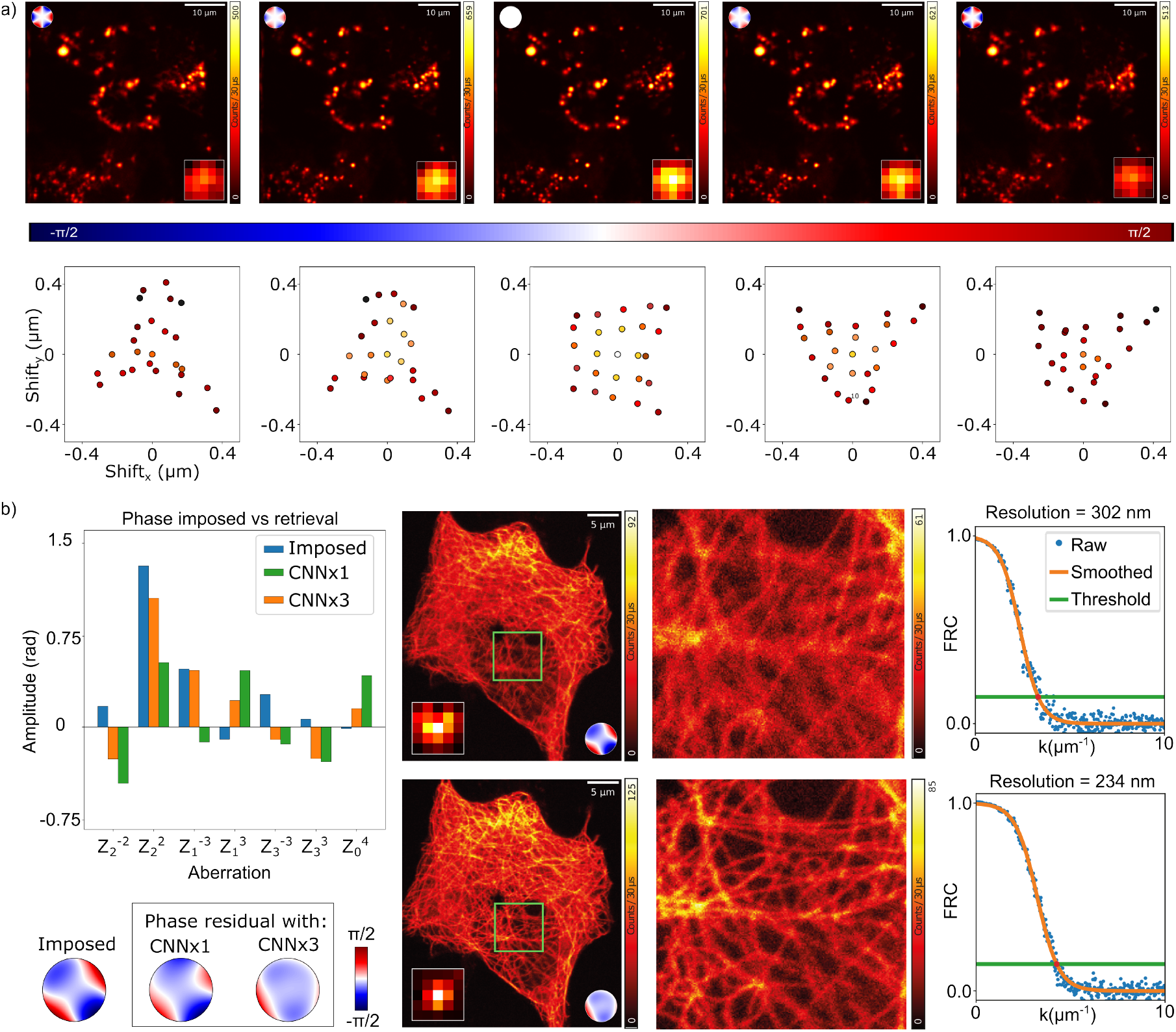
The panel a) illustrates the data acquisition process for transfer learning. In this instance, a specific aberration (trefoil 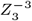) is employed, with an amplitude range of 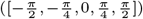. The top left corner of the image displays the relative phase mask applied to the deformable mirror. In the bottom right, the relative fingerprint obtained from the detector array is shown. Subsequently, data augmentation techniques are applied to enhance the dataset by increasing the number of experimental images. In the first row, the image with an imposed aberration is presented alongside a zoomed-in section of the cells highlighted by the blue square. The resulting image exhibits noticeable blurring, and the structures within the sample are poorly defined. In the second row -panel b-is reported an experimental dataset with fixed cells, which presents 3D structures not well defined due to the presence of random optical aberrations. The same image is showcased after correction as predicted by CNN, which reveals the presence of astigmatism as the main aberration of the system. Even if the prediction is not perfectly aligned with the imposed aberration, there is an improvement in image resolution and SNR compared to the previous iteration. Tubulin filaments, in particular, become more distinguishable. Concluding with the last row, the ground truth, an image of the same cell under aberration-free conditions.

The newly trained CNNs were subsequently tested on an entirely novel dataset obtained by imaging the cytoskeleton of a cell. This dataset represented a true three-dimensional sample, containing both in-focus and out-of-focus structures, thereby providing a realistic and challenging validation scenario. To validate the ISMAO approach, we introduced three aberrations simultaneously during the imaging experiments, each with random amplitudes, resulting in a RMSE of 1.4 rad (Fig. 5 b). The resulting image displayed noticeable blurring, making tubulin filaments indistinguishable, particularly those highlighted in the green box. The calculation of the FRC on the raw image estimated a resolution of 302 nm – well above the expected resolution. Subsequently, correlograms were computed from the aberrated image, and the CNN estimated the imposed aberrations, applying corrections that yielded the improved image, where the tubulin structures became visible compared to the previous one. This correction also led to an increase in SNR of 24% and resolution of almost 76%. This enhancement demonstrates the ability of the CNN and the ISMAO approach to discern intricate details within complex cellular structures.

## Conclusion

We introduced ISMAO, a novel sensorless aberration correction method for ISM that leverages a custom-designed CNN to decode the intrinsic optical aberration information encoded within LSM imaging datasets acquired using a detector array. Through pre-processing steps applied to the ISM imaging dataset – grounded in the image formation principles of ISM and designed to provide the CNN with more informative inputs – our approach achieves robust and efficient aberration prediction. Furthermore, the pre-processing steps, particularly the calculation of the cross-correlogram, render the method independent of sample structure, making ISMAO broadly applicable.

The CNN of ISMAO is trained on synthetic ISM datasets generated through a comprehensive simulation framework and achieves high accuracy in predicting aberrations under varied imaging conditions. Furthermore, by incorporating transfer learning, the method leverages pre-trained knowledge to enhance performance, even with limited experimental data. A key innovation of ISMAO is its ability to deliver aberration predictions from a single focal-plane image, with further improvements observed when using a multi-plane acquisition strategy. With just three planes, this approach enriches the network’s input with diverse and informative data, significantly improving its capacity to decode complex aberration patterns.

The core principle of the ISMAO approach lies in the ability of detector array imaging to capture information about the LSM point spread function (PSF) and specifically the optical aberrations influencing the imaging experiment. Building on this, we provide an alternative method for decoding optical aberrations from the imaging dataset. For example, more conventional regression analysis could be applied to the fingerprint, especially as larger detector arrays (e.g., 7 × 7 elements) become available, potentially offering greater precision and flexibility.

The advancements of this project will not only improve realtime adaptive optics imaging through the fast predictive capabilities of ISMAO to estimate optical aberration but also facilitate the development of sophisticated post-processing algorithms, such as multi-image deconvolution with realistic PSFs directly estimated from the imaging dataset. By further refining this framework, we aim to unlock new possibilities in next-generation microscopy.

We have demonstrated the advantages of ISMAO on a custom ISM setup equipped with a SPAD array detector. The unique spatiotemporal information provided by the SPAD array enables seamless integration of ISMAO benefits with other advanced microscopy techniques. These include stimulated-emission depletion microscopy (STED) (40, 42, 45), fluorescence correlation spectroscopy (FCS) (49, 50), and fluorescence lifetime imaging (FLIM) (51–54). Additionally, SPAD-based ISM implementations have been successfully combined with two-photon excitation (55) and, more recently, with single-molecule localization microscopy (56, 57). These results open up exciting opportunities for followup research aimed at integrating ISMAO (58) into these advanced microscopy techniques.

## Methods and materials

### Optical architecture

The experimental setup for ISM comprises a deformable mirror (ALPAO97, Alpao) and a single-photon-avalanche diode (SPAD) detector array (36) (Supp. Fig. S1). A continuous-wave laser with a wavelength of λ = 488 nm serves as the excitation source, coupled with a 4 mm lens (LA4024-A, Thorlabs) into a single-mode polarization-maintaining fiber (P5-405BPM-FC-2,Thorlabs). The light exiting from the fiber is collimated by a 50 mm lens (L2, LA1131-AB-ML, Thorlabs). A dichroic mirror (25 × 36 mm Longpass Dichroic Mirror, 490 nm cut-off) reflects the excitation laser, allowing fluorescence from the sample to pass through. The beam with a diameter of 15 mm reaches the DM (DM97-15 BAX255, pupil diameter 13.5 mm). The DM, positioned in the common path (excitation and fluorescence path), thus correcting aberrations in both illumination and detection paths. The laser beam then passes through lens pairs (L3 and L4 with focal lengths of 600 mm and 75 mm, respectively), producing an 8 × de-magnification of the excitation beam. The collimated beam is directed into the sample plane by galvanometric scanner mirrors XY (GVS102, Thorlabs). The pivot point of the scanner is projected by the scan lens (SL, focal length: 50 mm, Leica Microsystems) and a tube lens (TL, focal length: 200 mm, Leica Microsystems) into the back aperture of the objective lens (HCX PL APO 100 × */*1.4 −0.7 Oil CS, Leica Microsystems) with a 5.6 mm diameter. To ensure overfilling, the pair of lenses SL and TL produces a 4 magnification of the excitation beam spot onto the objective pupil. Although the pupil of the DM measures 13.5 mm in diameter, after the de-magnification the objective back aperture should be overfilled at least for 10% of its dimension. Otherwise, the contribution of the more lateral actuators can be lost. Then, the pair of lenses (SL and TL) produce a magnification of 4 × of the excitation beam spot onto the objective pupil. The fluorescence signal (λ = 510 − 520 nm) is collected by the objective lens and de-scanned by the galvanometric mirrors, filtered by DM1, and directed through lenses (L5 with a focal length of 600 mm, L6 with a focal length of 50 mm, LA1422-A-MLd, Thorlabs, and L7 with a focal length of 150 mm, LA1433-A-MLd, Thorlabs) to the detector array. The final lens (L7) images the fluorescent spot onto the detector, with a total magnification of M = 450. Our BrightEyes-MCS software acquires the raw data and control all part of the microscope (59).

### Deformable mirror calibration

After integrating the deformable mirror (DM) into the optical path of the LSM setup, we calibrated it to achieve the desired wavefront modulation. The calibration involved intercepting the excitation beam after the DM using a removable mirror and directing it through a telescope system onto a Shack-Hartmann sensor (SHS, SHCMOS, Alpao). The telescope was designed to conjugate the DM plane to the microlenses of the SHS and to provide a magnification that aligns the DM size with the SHS active area. The true calibration is finally obtained by using the ALPAO Core Engine (ACE). Additionally, we used the same SHS to correct aberrations potentially introduced by the optical elements between the DM and the objective lens. For this purpose, we introduced a beam splitter before the objective lens to direct the excitation beam onto the SHS. Also in this case, a telescope was used to conjugate the SHS plane and match the beam size effectively.

### Synthetic images for training

The generation of the synthetic dataset for training the CNNs is based on the BrightEyes-ISM package (46). This package not only offers various reconstruction algorithms for ISM dataset but also provides tools for generating physics-based simulated ISM scanned images. Specifically, it includes: (i) functions for creating three-dimensional phantom structures that combine point-like features with more complex filamentous structures both parametrized (e.g., size, density); (ii) a point-spread function generator based on vectorial modeling of the focused electromagnetic field, capable of accounting for the detector array geometry, various imaging conditions, and integrating any optical aberrations; and (iii) a simulator for generating scanned ISM datasets by convolving phantom structures with PSFs and incorporating different noise sources (e.g., photon counting noise, thermal dark noise). Since each element of the detector array generates a distinct scanned image, a 2*D* PSF was calculated for each element. To ensure realistic PSFs, the simulations were parameterized using the optical and physical properties of the microscope and detector array. Specifically, the excitation wavelength was set to 488 nm, while the emission wavelength was centered at 515 nm with a bandwidth of ±5 nm. The oil refractive index was assumed to be 1.5, and the numerical aperture (NA) of the objective was 1.4.

A pixel size of 50 nm and a magnification of 450 × at the detector array plane were used. The detector was modeled as a 5 × 5 square array with a pixel pitch of 75 *µ*m and a pixel size of 50 *µ*m.

To calculate the out-of-focus PSFs for generating the CNNx3 training set, a depth of field of 390 nm was assumed. Optical aberrations were introduced into the PSFs by modeling the wavefront at the pupil of the objective lens using a Zernike polynomial basis (with polynomials ranging from 5^*th*^ to 11^*th*^).

To randomize the aberrations in the training dataset, the coefficients of the Zernike polynomials were sampled from a double Gaussian distribution with a mean of *µ* = ±0.75 rad and a standard deviation of *σ* = 0.3 rad. The coefficients were then constrained in modulus to values between 0.25 r to 1.5 rad. Additionally, the 2*D* structures generating the images were randomized to increase diversity. A mixture of point-source structures and tubular structures was used, with the paths and sizes of the tubular structures generated randomly. Their concentrations, specifically their emission flux, were also randomized to capture a wide range of potential imaging scenarios. Given the PSFs and the phantoms, the images are generate with a simple 2D convolution. Two sources of noise are successively introduced to the images. The choice of a random pixel-dwell time between 30-60 *µs* for each simulation combined with the different photon-flux of the phantom allows the generation of images with different photon-counting noise. Furthermore, we also introduce the dark-noise intrinsic to each element of the SPAD array detector. A realistic value of dark-noise for the array element is measured directly from the SPAD array used in the custom ISM setup. A comprehensive table detailing all parameters used to generate the synthetic images is available in Suppl. Tab. S1. To train the custom CNNs, we generated a dataset comprising a total of 10,000 ISM experiments. This dataset had dimensions of 10,000 (phantoms) × 25 (array elements) × 3 (axial planes), amounting to nearly 110 GB of data. The simulations were performed on the Franklin supercomputer provided by the Istituto Italiano di Tecnologia (IIT) and required approximately one week to complete. After calculating the normalized correlograms, only the central region, typically consisting of 128 × 128 pixels was used for training the CNN. This choice was motivated by the fact that the central region contains most of the relevant information about the aberration, whereas the peripheral regions are dominated by noise.

### CNN architecture

Here, we describe the CNN3 architecture, as CNN1 can be considered its simplified version, reduced to process only the images of the in-focus plane. CNN is inspired by AlexNet (60, 61), and consists of three parallel branches, each designed to process the same input shape of (24, 128, 128) × 3 (Supp. Fig. S9). In each branch, the first step is a convolutional layer that applies 16 filters of size 5 × 5 to extract initial features from the input. This is followed by batch normalization, which normalizes the outputs to stabilize the learning process and improve convergence. Next, a max pooling layer reduces the spatial dimensions of the feature maps by taking the maximum value in each 2 × 2 block, helping to decrease the number of parameters and computational load. The branch continues with another convolutional layer, this time applying 32 filters of size 3 × 3, followed by batch normalization and another max pooling layer. This process is repeated with a third convolutional layer that applies 64 filters of size 2 × 2, once again followed by batch normalization and max pooling. At the end of each branch, the output is flattened into a one-dimensional vector, preparing it for the subsequent fully connected layers. After all three branches have processed the input, their outputs are concatenated, merging the distinct features learned from each path. This combined representation is then fed into a dense layer with 1024 units, allowing for complex feature combinations. To prevent overfitting, a dropout layer randomly drops 50% of the units during training. This is followed by another dense layer of 1024 units and another dropout layer. Finally, the network concludes with a dense layer that outputs a vector of size 7, which likely represents class scores for a multi-class classification task. Overall, this architecture effectively combines feature extraction and classification, making it suitable for various image-processing applications. To assess the effectiveness of our CNN models in predicting Zernike polynomial aberrations, it is crucial to employ appropriate evaluation metrics that provide meaningful insights into both accuracy and precision. Given the regression nature of this problem, we focus on metrics that quantify the error or correlation between the predicted and true labels corresponding to the aberration amplitude. The Root Mean Square Error (RMSE) is chosen as the loss function over the amplitude of the Zernike polynomials, with 300 epochs and a batch size of 64. This configuration takes approximately 1.5 hours for CNNx1 and 4 hours for CNNx3. For the transfer learning process, we used only 30 epochs since the model begins with weights that are already close to optimal, and the training process tends to converge more quickly. This faster convergence reduces the need for extensive training over many epochs.

**Table 1.** Numbers of parameters of CNNx1 model for each layer.

| Layer | Type | Output Shape | Param # |
| --- | --- | --- | --- |
| 1 | Conv2D | (42, 42, 16) | 9616 |
|  | BatchNorm | (42, 42, 16) | 64 |
|  | MaxPool2D | (21, 21, 16) | 0 |
| 2 | Conv2D | (10, 10, 32) | 4640 |
|  | BatchNorm | (10, 10, 32) | 128 |
|  | MaxPool2D | (5, 5, 32) | 0 |
| 3 | Conv2D | (4, 4, 64) | 8256 |
|  | BatchNorm | (4, 4, 64) | 256 |
|  | MaxPool2D | (2, 2, 64) | 0 |
| 4 | Dense | 256 | 787456 |
|  | Dropout | 0.5 | 0 |
| 5 | Dense | 1024 | 1049600 |
|  | Dropout | 0.5 | 0 |
| 6 | Dense | 7 | 7175 |

### Transfer learing

Training a CNN from scratch requires large datasets and significant computational resources. By starting with a pre-trained model, which already captures general features from a related domain, transfer learning fine-tunes the network for the specific task, requiring fewer epochs and less data in order to improve the generalization capabilities of CNNs. This strategy is particularly beneficial in contexts where labeled data is limited. By utilizing experimental data obtained from observations of fluorescence beads and lipid droplets, we captured features related to real-content conditions, which are critical to replicate in simulations. We collected nearly one hundred experimental images for each focal plane. A subset of this dataset is presented in Suppl. Fig. S8 showcasing experimental images of beads under various amplitudes of trefoil 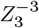 aberration. Given the time-intensive process of acquiring thousands of experimental images for CNN training, we employed transfer learning combined with data augmentation techniques on the collected experimental dataset. To further enhance the dataset, we generated an additional 1000 images by cropping the original images around randomly selected central points. To ensure the quality of the augmented data, we applied a threshold to exclude entirely dark images that might result from cropping. However, due to the nature of the dataset, the data augmentation techniques were limited only to the cropping function; we could not perform all possible geometric transformations on the experimental images, as this would distort the correlograms and affect the classification of the aberrations. Overall, the transfer learning approach aims to enhance the CNNs capability to detect and estimate optical aberrations effectively in practical applications, for this reason, the number of images in the experimental dataset post-augmentation constituted approximately only 10% of the overall data used to train the CNN taking almost only 5 minutes to complete the process.

### Image processing

#### Confocal image

Given the ISM image dataset (Eq. 1), we generated the corresponding confocal image by summing all the raw images:

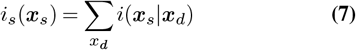

The result is equivalent to a confocal image acquired with a pinhole as large as the detector array.

#### Shift-vectors and adaptive pixel reassignment

We calculated the shift-vectors of an ISM dataset as:

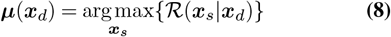

The APR reconstruction is calculated as the sum of the aligned images

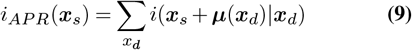

#### Correlograms

The correlograms are calculated from an ISM dataset as follows. The cross-correlation between an image of the ISM dataset and the reference image at the center of the array detector (*x*_*d*_ = 0) is calculated as:

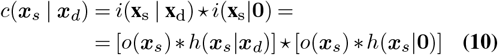

where * and * are the convolution and the cross-correlation operator, respectively. Then, we apply the Fourier transformation to the scan coordinates *x*_*s*_ obtaining:

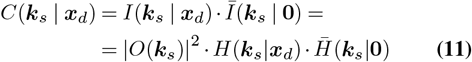

where the overline represents the complex conjugate operation, *k*_*s*_ is the scan spatial frequency, and the uppercase letters indicate the Fourier transform of the quantities previously indicated with the lowercase letter. To discard the object contribution, we need to normalize the correlation in Fourier space.

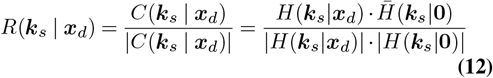

Note that the denominator of the above division might contain zeros of very small numbers. To avoid divergence, we impose the correlogram to be zero in the positions where the denominator value is smaller than the threshold of *ε* = 1 × 10^−7^.

We normalized by the fingerprint, defined as follows

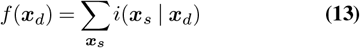

Finally, we obtain the correlograms by inverting the Fourier transform and we normalize with the fingerprint

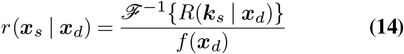

#### Sample preparation

To produce a sample with fixed fluorescent beads we first deposit 150*µL* of poly-L-lysine (PLL) on a clean cover slip and incubate it at 37°C for 10 minutes. In the meanwhile, we prepare a dilution of the beads’ mother solution in distilled water with a volumetric ratio between 1 : 500 and 1 : 500, that we sonicate for 5 minutes. We dry the coverslip with clean air and deposit 150*µL* of beads dilution on top of the adhesive film, followed by incubation at at 37°C for 10 minutes. We spill the remainder of the solution on the cover glass and dry with clean air. We add 5*µL* of Mowiol^®^ mounting medium and seal the cover glass on a microscope slide.

HEPG2, a human hepatocellular carcinoma cell line, and HeLa, a widely known cervical carcinoma cell line, were maintained respectively in Eagle’s Minimum Essential Medium (EMEM) and Dulbecco’s modified Eagle medium (D-MEM), both containing 10% Foetal Bovine Serum (FBS), at 37°C. For imaging, cells were seeded on poly-Llysine coated glass slides. When cells reached the proper confluence, slides were rinsed with PBS, fixed with 4% paraformaldehyde/PBS for 15 minutes, then washed again three times with PBS.

Lipid droplets (LDs) in HEPG2 cells were visualized using Nile Red, a fluorescent dye with a high sensitivity to neutral lipids. After fixation and washing, HEPG2 cells were incubated with 0.5 ug/ml Nile Red (Invitrogen, Thermo Fisher Scientific) in PBS for 30 minutes, washed with PBS and mounted using ProLong™ Diamond Antifade Mountant (Thermo Fisher Scientific). Beta-tubulin staining was performed on HeLa cells. To permeabilize cells and block the non-specific binding of the antibodies during the immunostaining, we incubated the coverslides with 3% BSA in 0.5% Triton *X* − 100 in PBS for 1 h, at room temperature. Slides were then incubated with 1 : 200 rabbit anti-b tubulin antibody (ab6046, Abcam) overnight at 4°C. The following day, after washing three times with 0.5% Triton-PBS, 1 : 500 Alexa Fluor 488 anti-rabbit antibody (*a*11008, Abcam) was used as secondary antibody and incubated at room temperature for 1 h. Then, slides were rinsed and mounted as above.

## ACKNOWLEDGEMENTS

This research was supported by: the European Research Council, BrightEyes No. 818699 (G.V.). The authors thank all members of the Molecular Microscopy and Spectroscopy labs for the many helpful suggestions: Sabrina Zappone, Dr. Eleonora Perego, Dr. Mattia Donato, Dr. Eli Slenders, Giacomo Garré, Dr. Andrea Bucci, Luca Bega, Sanket Patil, Dr. Sami Walteri Koho, and Dr. Marcus Held.

## AUTHOR CONTRIBUTIONS

G.V. conceived the idea. G.V. designed the study. A.Z., and G.V. supervised the project. F.F. built the microscope with the adaptive optics element. M.D. implemented the data acquisition and control system. F.F. and F.B. designed the cell experiments. F.F. developed the convolutional neural network with the supervision of P.M. F.F. analyzed the data with the support of all other authors. F.F. and G.V. wrote the manuscript. All authors discussed the results and commented on the manuscript.

## CODE

The Python code used in this work is available at the following GitHub repository: https://github.com/VicidominiLab/ISMAO. The experimental data generated for this study are available in the following Zenodo database: https://doi.org/10.5281/zenodo.13789465.

## COMPETING INTERESTS

G.V. has a personal financial interest (co-founder) in Genoa Instruments, Italy.

## Notes

https://github.com/VicidominiLab/ISMAO

https://doi.org/10.5281/zenodo.13789465

