## Supplementary Figures for "Wavefront estimation through structured detection in laser scanning microscopy"

### Supplementary Note 1:

**ISMAO microscope**

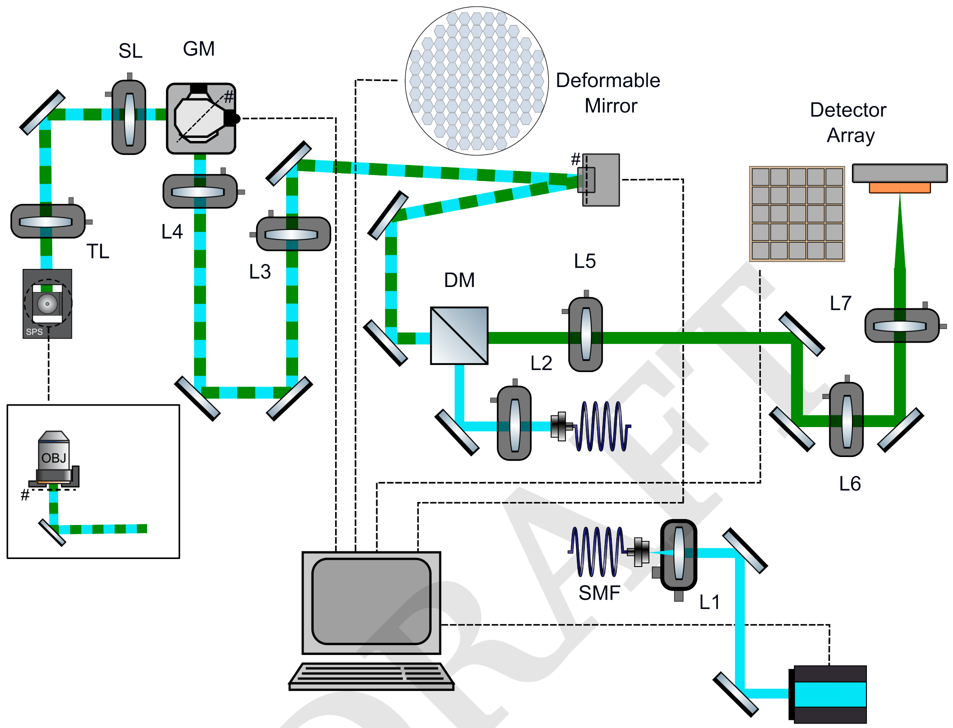


DRAFT

**Fig. S1.** The experimental setup for ISM with AOE comprises an Alpao 97 actuators deformable mirror and a dedicated detector array. Laser with a wavelength of $\lambda = 488$ nm. L1 = $4$ mm, L2 = $50$ mm, L3 -L5 0 $600$ mm, L4 = $75$ mm and L6 =$50$, L7 = $150$ mm . Single-Mode polarization-maintaining Fiber (SMF). Dichroic mirror longpass (DM), Scan Lens (SL = $50$ mm,) and a Tube Lens (TL= $200$ mm). Objective Lens (OBJ Leica HCX PL APO 100×1.4-0.7 Oil CS). DM (DM97-15 BAX255. Array detector $5 \times 5$.

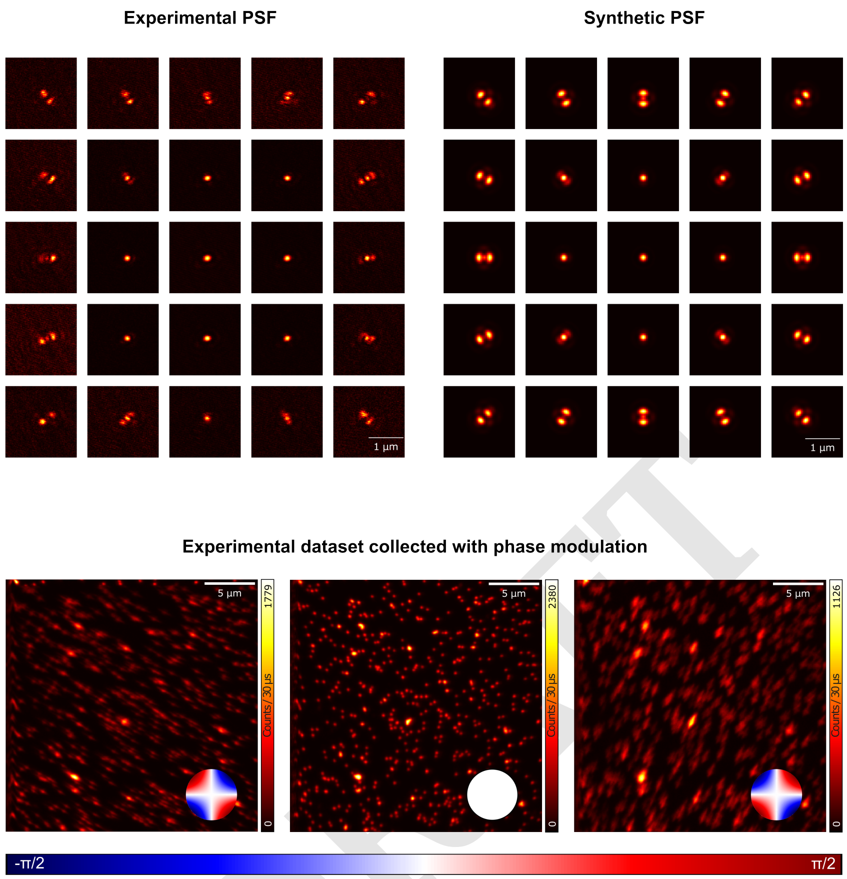


**Fig. S2.** A visual comparison between the real and simulated PSFs from the ISMAO setup is presented. The top image shows scattered light collected from $80$ nm beads, demonstrating the real PSF. The bottom image features fluorescence beads $(100$ nm) after introducing phase modulation using the DM following prior calibration. The phase modulation induces oblique astigmatism, which is visible in the imaging results.

| Parameter | Value |
| --- | --- |
| Ex. wavelength | $488\,\mathrm{nm}$ |
| Em. wavelength | $515\,\mathrm{nm}$ |
| NA | $1.2$ to $1.4$ |
| Ex. gamma | $45°$ |
| Ex. beta | $90°$ |
| Refractive index | $1.5$ |
| Phase mask sampling | $250$ |
| Zernike polynomial | $5$ to $11$ |
| Amplitude polynomial | $\pm\pi/2\,\mathrm{rad}$ |
| Pixel size | $50\,\mathrm{nm}$ |
| # lateral sampling | $61\,\mathrm{pixel}$ to $81\,\mathrm{pixel}$ |
| # axial sampling | $1$ to $3$ |
| Magnification | $450$ |
| Pixel pitch | $75\,\mathrm{\mu m}$ |
| Pixel size | $50\,\mathrm{\mu m}$ |
| # elements | $25$ |
| Depth of Field | $390\,\mathrm{nm}$ |
| Zernike $\mu$ | $0$ |
| Zernike $\sigma$ | $0.3$ |
| Sample | Tubulin, beads, or mix |
| Sample pixel | Pixel size |
| Beads diameter | $100\,\mathrm{nm}$ |
| Photon flux | $10\,\mathrm{MHz}$ to $40\,\mathrm{MHz}$ |
| Dwell time | $30\,\mathrm{\mu s}$ to $60\,\mathrm{\mu s}$ |
| Field of view | $201\,\mathrm{pixels}$ to $501\,\mathrm{pixels}$ |

**Table S1.** This table summarizes the key parameters used in the experimental setup, including excitation and emission wavelengths, numerical aperture (NA), sample details, and imaging conditions. The values listed provide essential information for the optical system configuration, such as refractive index, phase mask sampling, pixel size, lateral and axial sampling, as well as sample properties like bead diameter and photon flux. These parameters are used to simulate a dataset, which is then utilized for training the two CNNs.

DRAFT

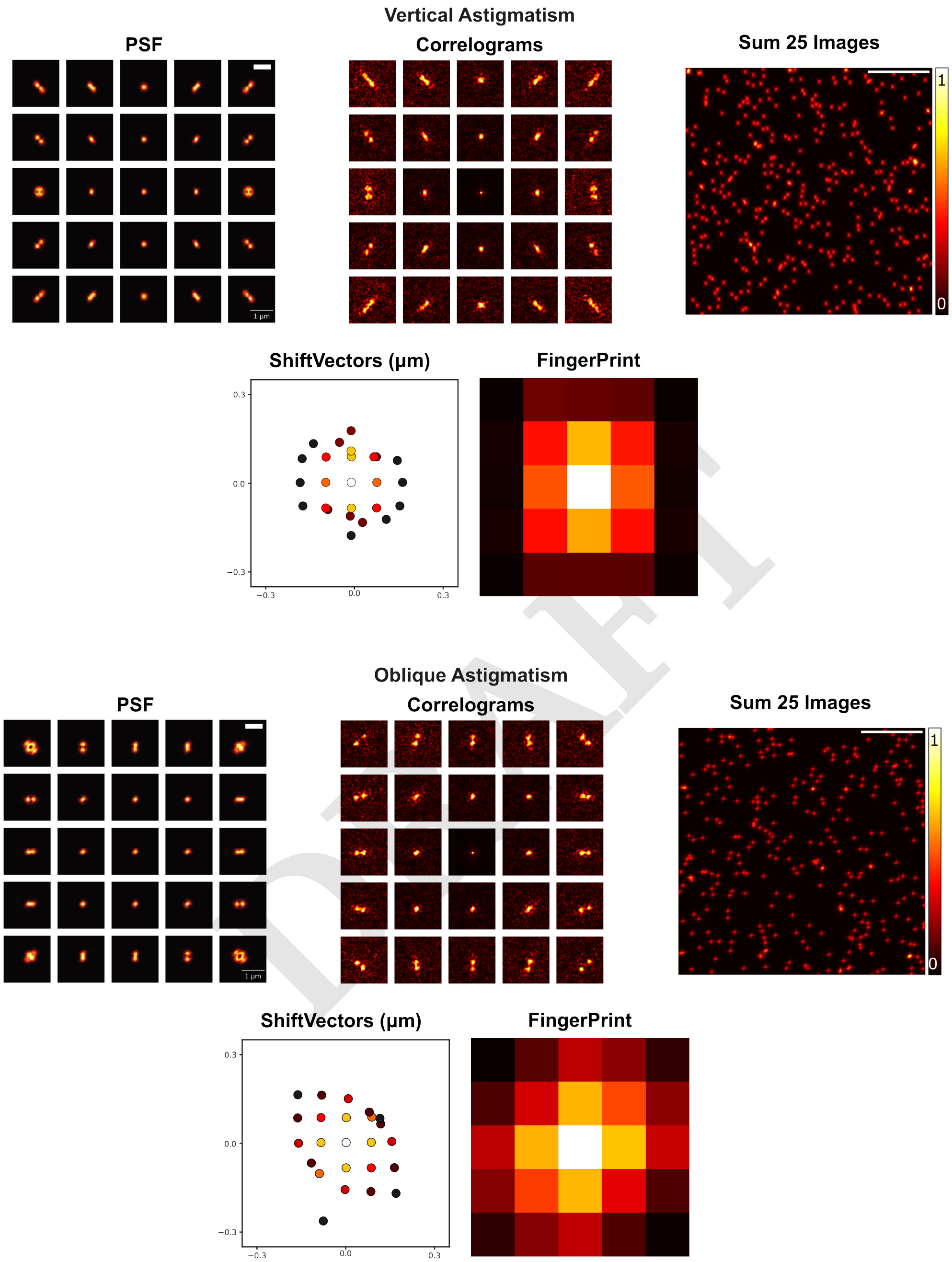


DRAFT

**Fig. S3.** Example of simulated PSFs, correlograms, sum of 25 scanned images, shift vectors, and fingerprint of *Vertical Astigmatism* and *Oblique Astigmatism* with an amplitude of $1$ rad. Astigmatism occurs when light rays entering the system are focused at different points along different axes, typically the horizontal and vertical planes, leading to blurred images, particularly in the axial direction. This results in the sample appearing elongated or elliptical instead of sharp. This effect is qualitatively reflected in the analysis of the shift vectors and fingerprint. Scalebar $1\mu m$ for PSFs images. Scalebar $10\mu m$ for the sum of 25 images.

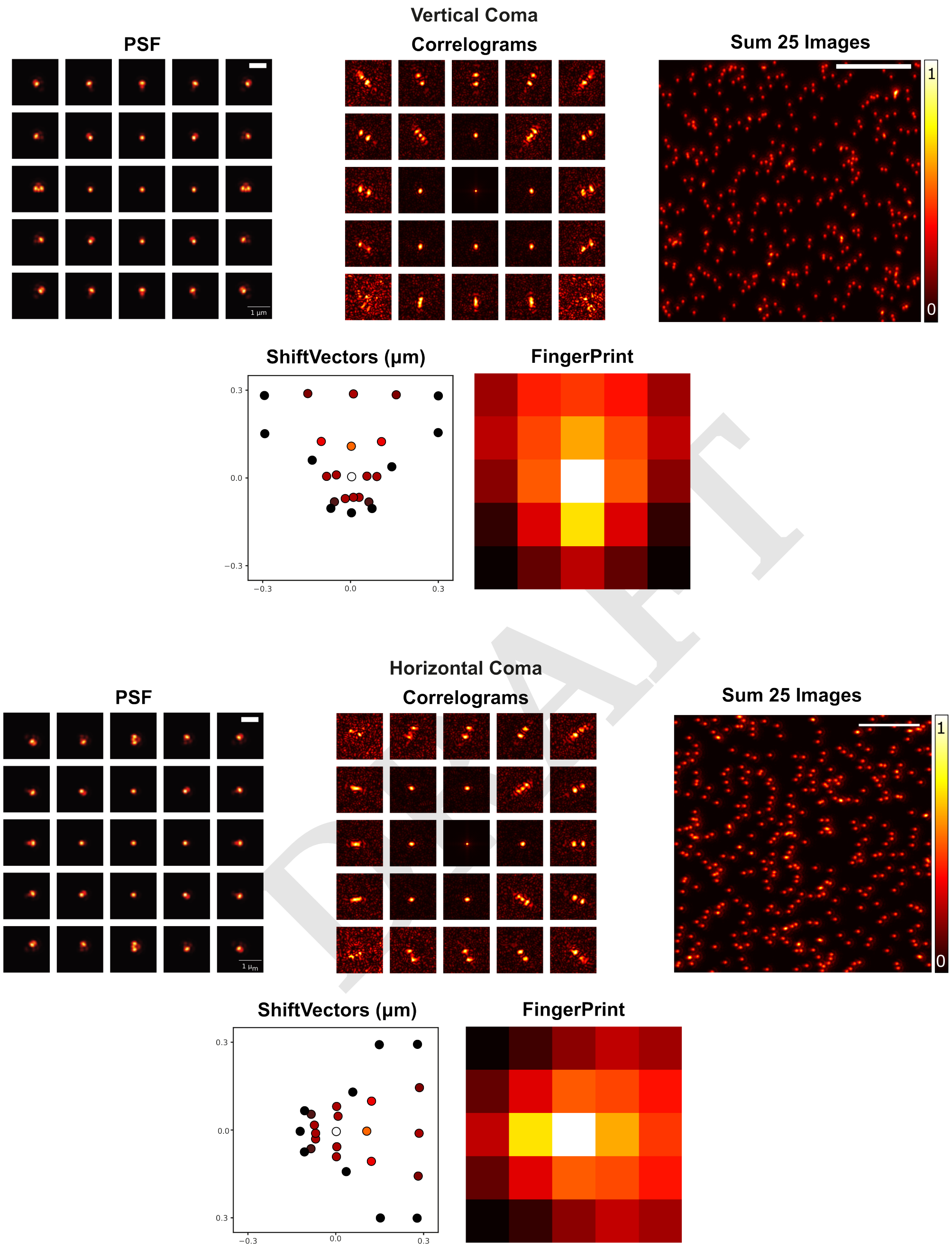


DRAFT

**Fig. S4.** Example of simulated PSFs, correlograms, sum of 25 scanned images, shift vectors, and fingerprint of *Vertical Coma* and *Horizontal Coma* with an amplitude of $1$ rad. Coma occurs when light rays passing through a system are unevenly focused, leading to asymmetric blurring, often resembling a comet-like shape. In the imaging, this distorts the shape of the sample, making it appear as tails, especially at the periphery of the field of view. A similar comet-like effect is also evident in the fingerprint along the direction of the aberration, as well as in the shift vectors. Scalebar $1\mu m$ for PSFs images. Scalebar $10\mu m$ for the sum of 25 images.

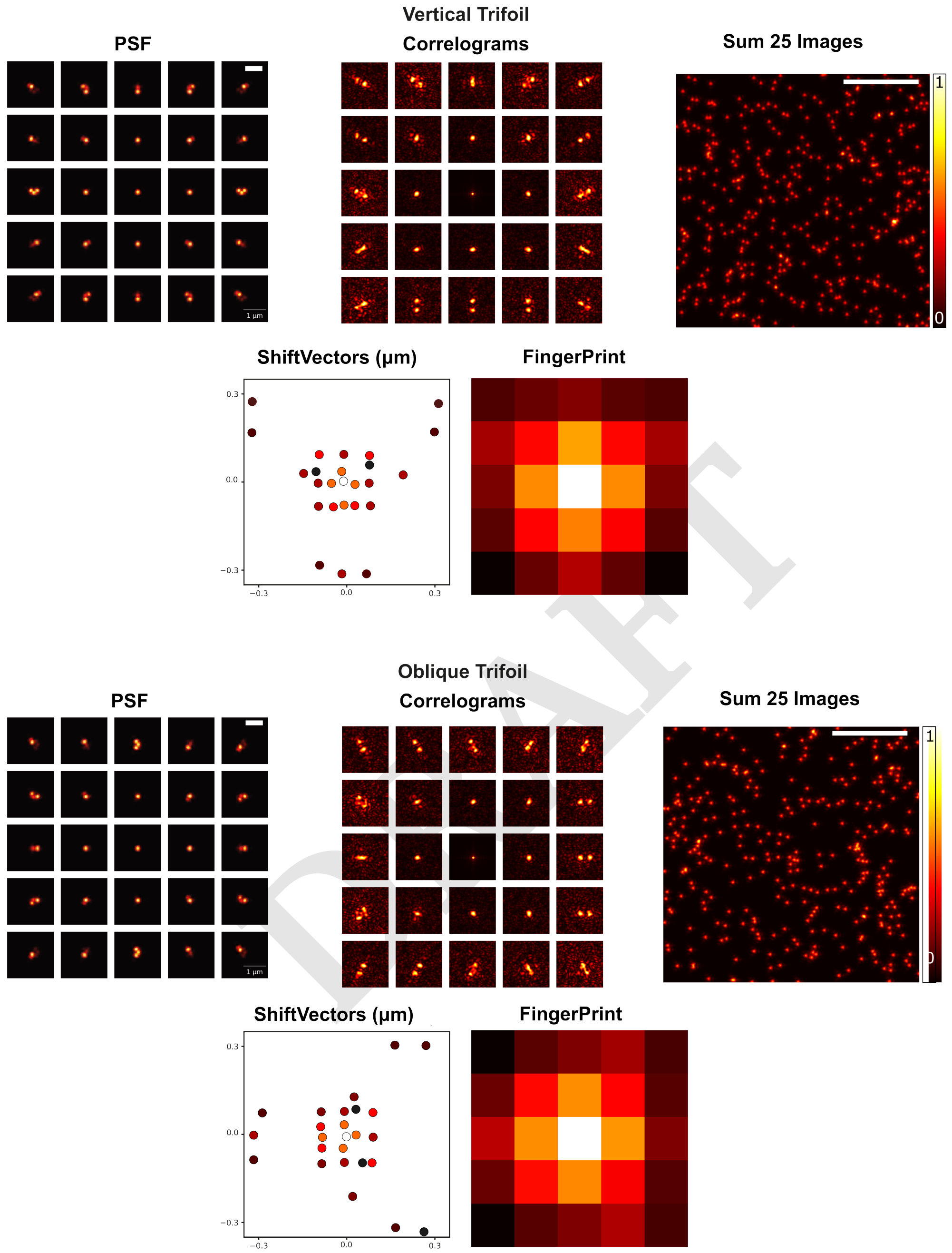


**Fig. S5.** Example of simulated PSFs, correlograms, sum of 25 scanned images, shift vectors, and fingerprint of *Vertical Trifoil* and *Oblique Trifoil* with an amplitude of $1$ rad. Trifoil is an optical aberration that occurs when light rays passing through a system are focused in three distinct directions, creating a pattern with three symmetrical lobes. Trifoil distorts the shape of beads, causing them to appear as a tri-lobed or triangular pattern rather than a sharp point. In this case, the structure of the aberrated PSF is also observed in the fingerprint and shift vectors, which follow the direction of the imposed aberration, resembling an arrow. Scalebar $1 \mu m$ for PSFs images. Scalebar $10 \mu m$ for the sum of 25 images.

DRAFT

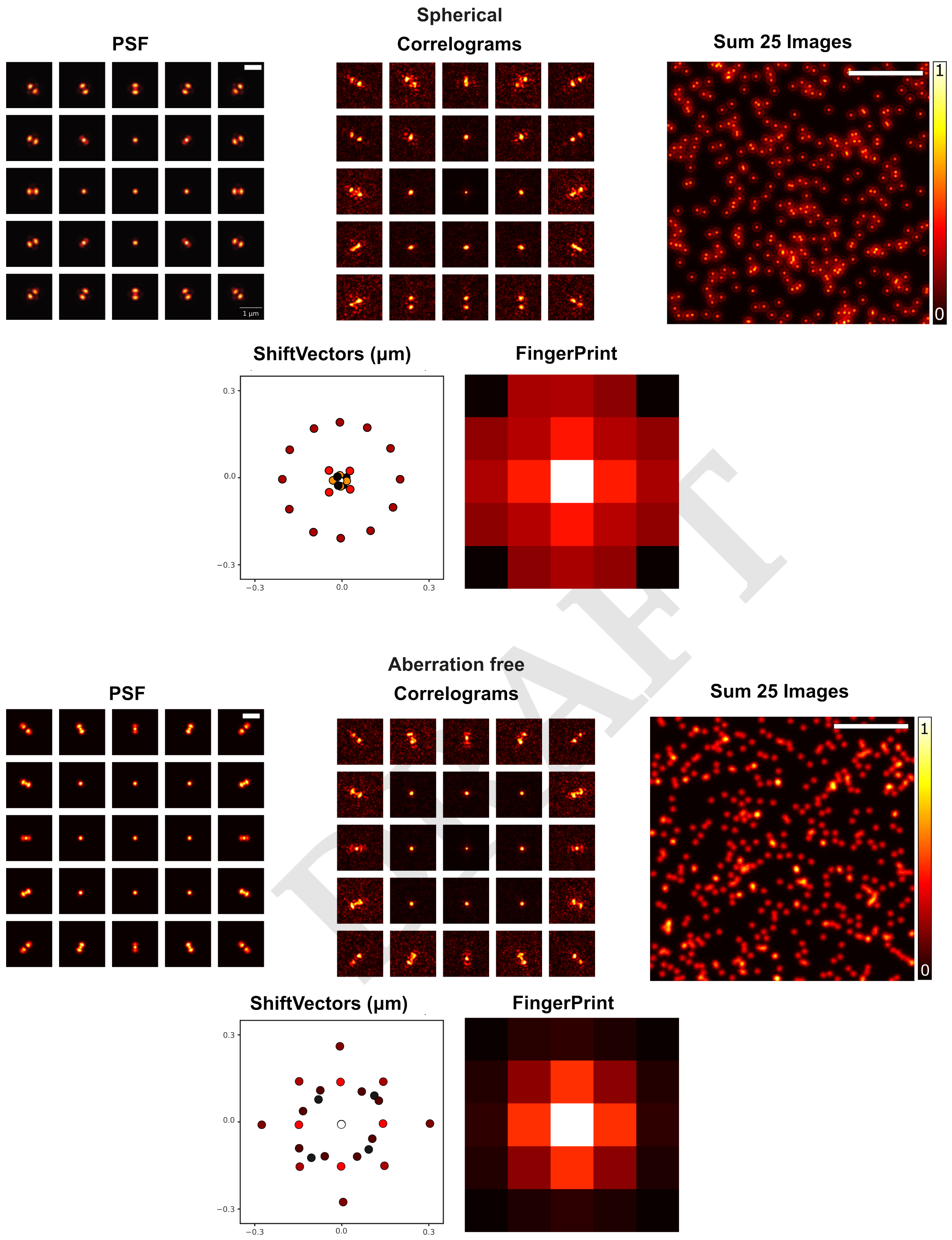


**Fig. S6.** Example of simulated PSFs, correlograms, sum of 25 scanned images, shift vectors, and fingerprint of *Primary Spherical Aberration* with an amplitude of $1\,\mathrm{r}$ to $1$ rad and *Aberration-Free.* Spherical aberration occurs when light rays passing through a lens or optical system are focused at different points along the optical axis, typically due to the curvature of the lens. The resulting image is blurred or unfocused. The imaging of beads appearing as distorted discs rather than sharp points. As for the previous aberration, also in this case can be immediatmy observed the distribution of the light alog the fingerprint which is pred along all the pixel of the sensor. Scalebar $1\mu m$ for PSFs images. Scalebar $10\mu m$ for the sum of 25 images.

DRAFT

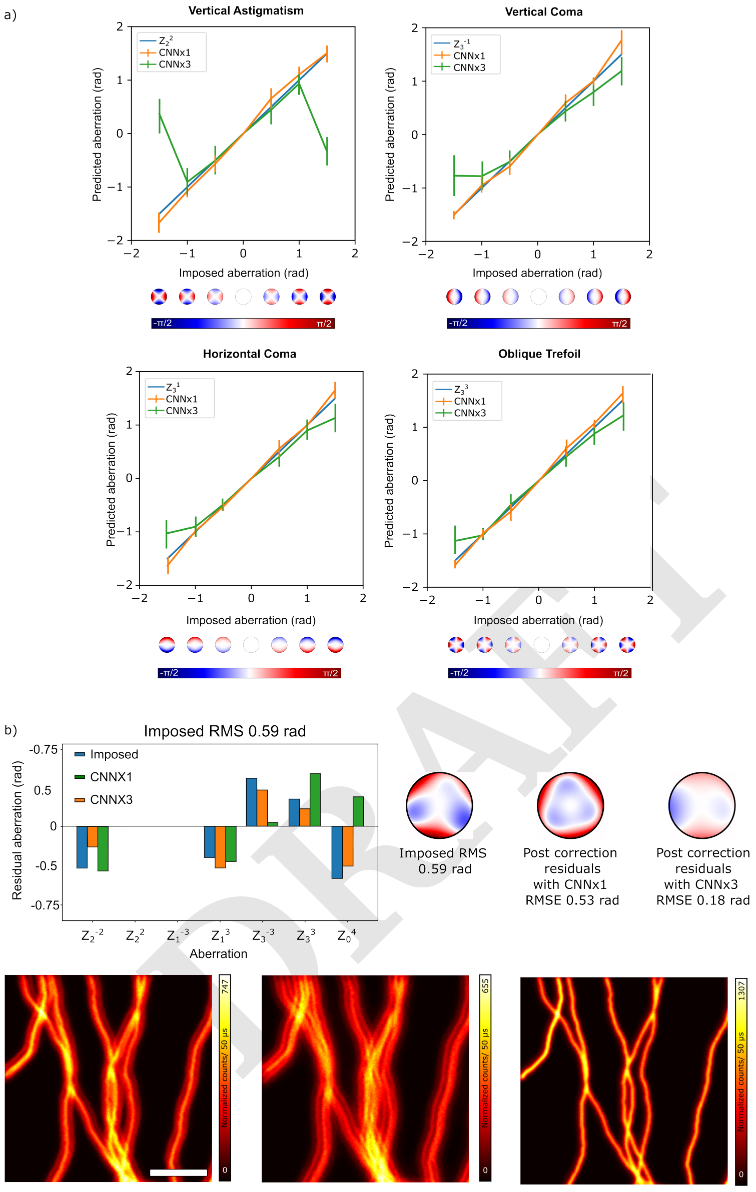


**Fig. S7.** a) Imposed aberrations and predictions. This panel presents a comparison of imposed aberrations, including Vertical Astigmatism, Vertical Coma, Horizontal Coma, and Oblique Trefoil. The amplitude range of the Zernike polynomials is indicated in blue ($-1.5$ and $1.5$ rad), while predictions from CNNx1 and CNNx3 are shown in green and orange, respectively. Each aberration is accompanied by its corresponding phase mask. The results demonstrate that CNNx1 effectively identifies aberrations with laterally varying correlograms, such as coma and trefoil. However, for aberrations with axial variation, such as vertical astigmatism, the predictive performance of CNNx1 deteriorates as the aberration amplitude increases, as discussed in the main text. b) Simulated Data at RMSE = 0.59 rad: In this scenario, the strongest imposed aberrations are Astigmatism and Spherical aberration. CNNx1 struggles to produce accurate predictions, generating outputs that further degrade image quality with an RMSE = 0.53 rad after the correction. Specifically, the large disparity between the imposed and predicted spherical aberration results in a phase pattern that fails to correct the image. In contrast, CNNx3 consistently delivers accurate predictions across all aberration types, even at higher amplitudes, maintaining excellent image quality with an RMSE = 0.18 rad after the correction.

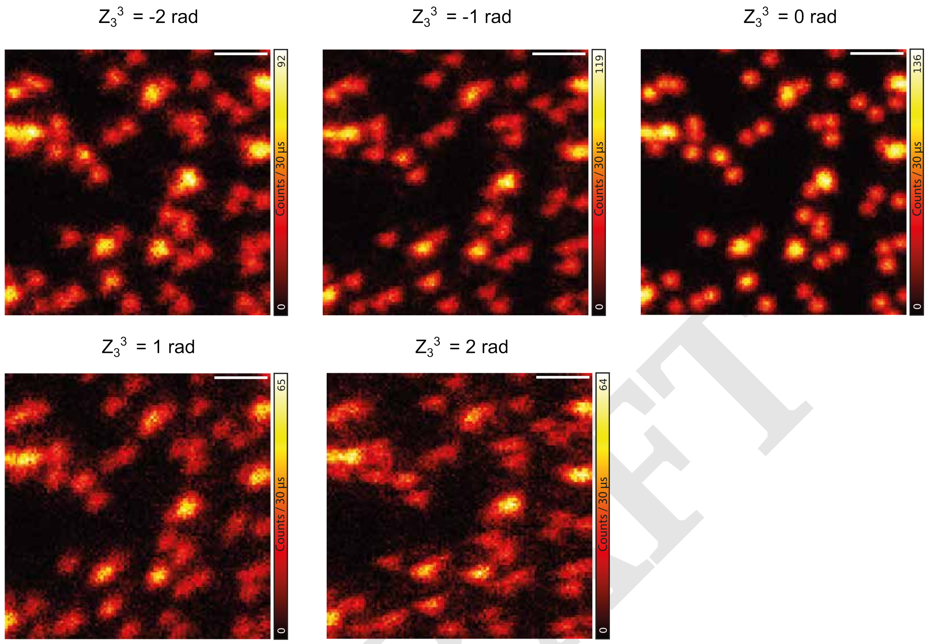


DRAFT

**Fig. S8.** Example of transfer learning, where fluorescence beads are used to collect aberrated images with an imposed trefoil aberration.

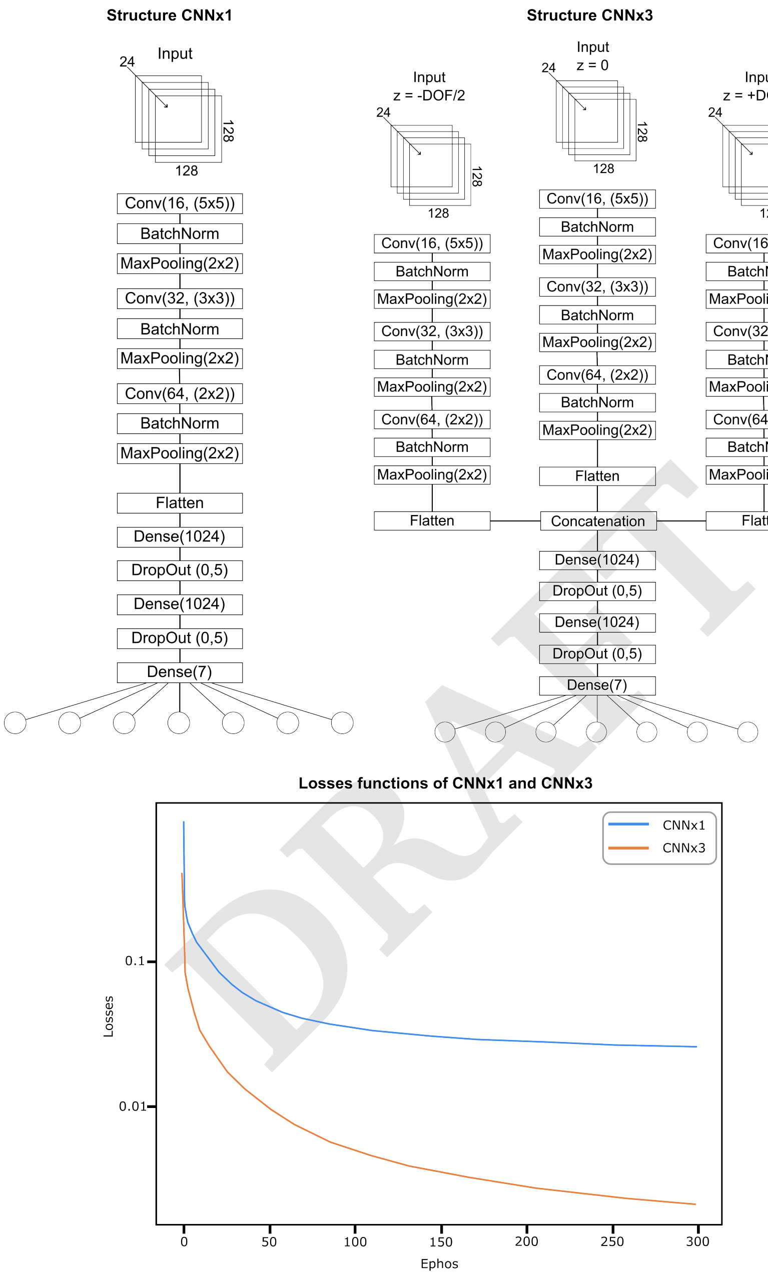


DRAFT

**Fig. S9.** At the top, the two architectures for CNNx1 and CNNx3 are illustrated. Both structures are similar, with the key difference lying in the input datasets. In the case of CNNx1, the input consists of a set of 24 correlograms generated from a single focal plane. In contrast, CNNx3 takes as input three layers of correlograms collected from different DOF. These three branches are concatenated after the flattening layer to combine the extracted features into a single branch, which is then used for outputs. At the bottom, the validation loss plot provides a graphical representation of the loss values calculated during the model's validation phase. The loss function quantifies the difference between the predicted and actual values, with CNNx1 yielding a loss of 0.3, while CNNx3 demonstrates significantly better performance with a loss of 0.01. This highlights the superior performance of CNNx3. The plot also illustrates the RMS loss function over epochs during the training of both architectures, using the same dataset but with different random splits for training and validation.
